# Investigating mechanisms of polarized light sensitivity in the small white butterfly *Pieris rapae*

**DOI:** 10.1101/2019.12.19.883272

**Authors:** Adam J. Blake, Gina S. Hahn, Hayley Grey, Shelby Kwok, Deby McIntosh, Gerhard Gries

## Abstract

There is an ever increasing number of arthropod taxa shown to have polarization sensitivity throughout their compound eyes. However, the mechanisms underlying arthropod perception of polarized reflections from objects such as plants are not well understood. The small white butterfly, *Pieris rapae*, has been demonstrated to exploit foliar polarized reflections, specifically the degree of linear polarization (*DoLP*), to recognize host plants. The well-described visual system of *P. rapae* includes several photoreceptor types (red, green, blue) that are sensitive to polarized light. Yet, the mechanism underlying the behavioral responses of *P. rapae* to stimuli with different *DoLPs* remains unknown. To investigate potential mechanisms, we designed several two-choice behavioral bioassays, displaying plant images on paired LCD monitors which allowed for independent control of polarization, color and intensity. We found that shifts in image intensity had a similar effect on *P. rapae* preferences for stimuli dissimilar in *DoLP* and dissimilar in color, suggesting *DoLP* differences are perceived as color. When a *DoLP* choice was offered between plant images manipulated in a manner to minimizing the response of blue, red, or blue and red photoreceptors, *P. rapae* shifted its preference for *DoLP*, suggesting a role for red, green and blue polarization-sensitive photoreceptors. Modeling of *P. rapae* photoreceptor responses to test stimuli suggests that differential *DoLP* is not perceived solely as a color difference. Our combined results suggest that *P. rapae* females process and interpret polarization reflections in a way different from that described for other polarization-sensitive taxa.

## Introduction

Polarized light cues are used by many arthropods but apart from polarized skylight navigation little is known about how these organisms perceive polarized reflections (Heinloth et al., 2018). All organisms with rhabdomeric photoreceptors have the potential to sense polarized light (Horváth and Varju, 2004). The photopigments within the membranes forming the microvilli of the rhabdom more strongly absorb light vibrating in a plane parallel to the microvillar direction. The direction of polarization, relative to the dorsal-ventral axis of the insect, giving a maximum photoreceptor response is referend to as *ϕ*_max_, whereas the ratio of response to light at *ϕ*_max_, and orthogonal to *ϕ*_max_, is referred to as polarization sensitivity (*PS*). Photoreceptors with a high *PS* are typically found in a specialized area of the compound eye known as the dorsal rim allowing for polarized skylight navigation (Labhart and Meyer, 1999). The microvilli of these photoreceptors are aligned, and non-twisted, along the length of their relatively short rhabdom, thereby enhancing *PS*. High *PS* can interfere with color vision, resulting in polarization-induced false colors (Wehner and Bernard, 1993). Many insects avoid these drawbacks by twisting the direction of these microvilli along the length of the rhabdom but many others, especially aquatic and semi-aquatic insects, possess photoreceptors with moderate *PS* throughout their compound eyes. Histological and electrophysiological work has also revealed evidence for *PS* in herbivorous insects.

Recently, *P. rapae* females have been shown to discriminate among potential host plants based on the polarization of light reflected from their foliage (Blake et al., 2019b). Like any shiny surface, the leaf surface preferentially reflects light oscillating parallel to that surface (Foster et al., 2018). This axis of polarization (*AoP*, 0-180°) is strongly dependent upon the viewing angle and the location of the light source but not on the characteristics of that surface (Blake et al., 2019b). While the *AoP* does not seem to carry useful host plant information, the degree to which the foliar reflection is polarized (degree of linear polarization, *DoLP*, 0-100%) does convey information about a leaf’s surface. As only the specular component of the reflection is polarized, any leaf traits that affects the relative shininess or mattness of leaves will alter the *DoLP*. Decreasing the diffuse reflection through absorbance by pigments, scattering the specular reflection with epicuticular waxes or pigments, or undulations of the plane of the leaf’s surface can all affect the *DoLP* of foliar reflections (Grant et al., 1993). Female *P. rapae* are able to discern cabbage host plants and potato non-host plants based on the lower *DoLP* of cabbage leaf reflections (Blake et al., 2019b). In choice bioassays, which presented manipulated host plant images, *P. rapae* females rejected most images with a *DoLP* dissimilar to that of their cabbage host plant. The informative value of this cue is enhanced by a relative insensitivity to most *AoPs* during host plant selection. Both the underlying neurological mechanism and the photoreceptors involved in this discrimination remain unknown.

The visual systems of *P. rapae* resembles that of other butterflies in that each ommatidium contains nine photoreceptors and the three ommatidial types are arranged in a random mosaic throughout the compound eye (Fig. 1a). Similar to the ommatidia of *Papilio* butterflies (Kelber, 2001), the shortwave (UV, violet, blue) sensitive R1,2 photoreceptors respond most strongly to polarized light, whereas the longwave-sensitive R3-9 photoreceptors respond most strongly to horizontally polarized light (R3,4) and obliquely polarized light (R5-8) (Blake et al., 2019a) (Fig. 2b,c). In the ventral portion of the eye, the sensitivity of the R5-8 photoreceptors are modified by perirhabdomal filtering pigments into red receptors distinct to each of three ommatidial classes, with more variation in *PS* among ommatidial types than reported in *Papilio* (Fig. 1c). Of the shortwave receptors, only the type I blue photoreceptors show significant *PS*. There is also a lower *PS* in type II R3,4 receptors, which may explain the difference in the axis of maximal polarization sensitivity (*ϕ*_max_) of red photoreceptors among ommatidial types. The R9 receptor is thought to be red-sensitive (Shimohigashi and Tominaga, 1991), and likely has low *PS* due to its bidirectional microvillar arrangement (Qiu et al., 2002).

**Fig. 1.**
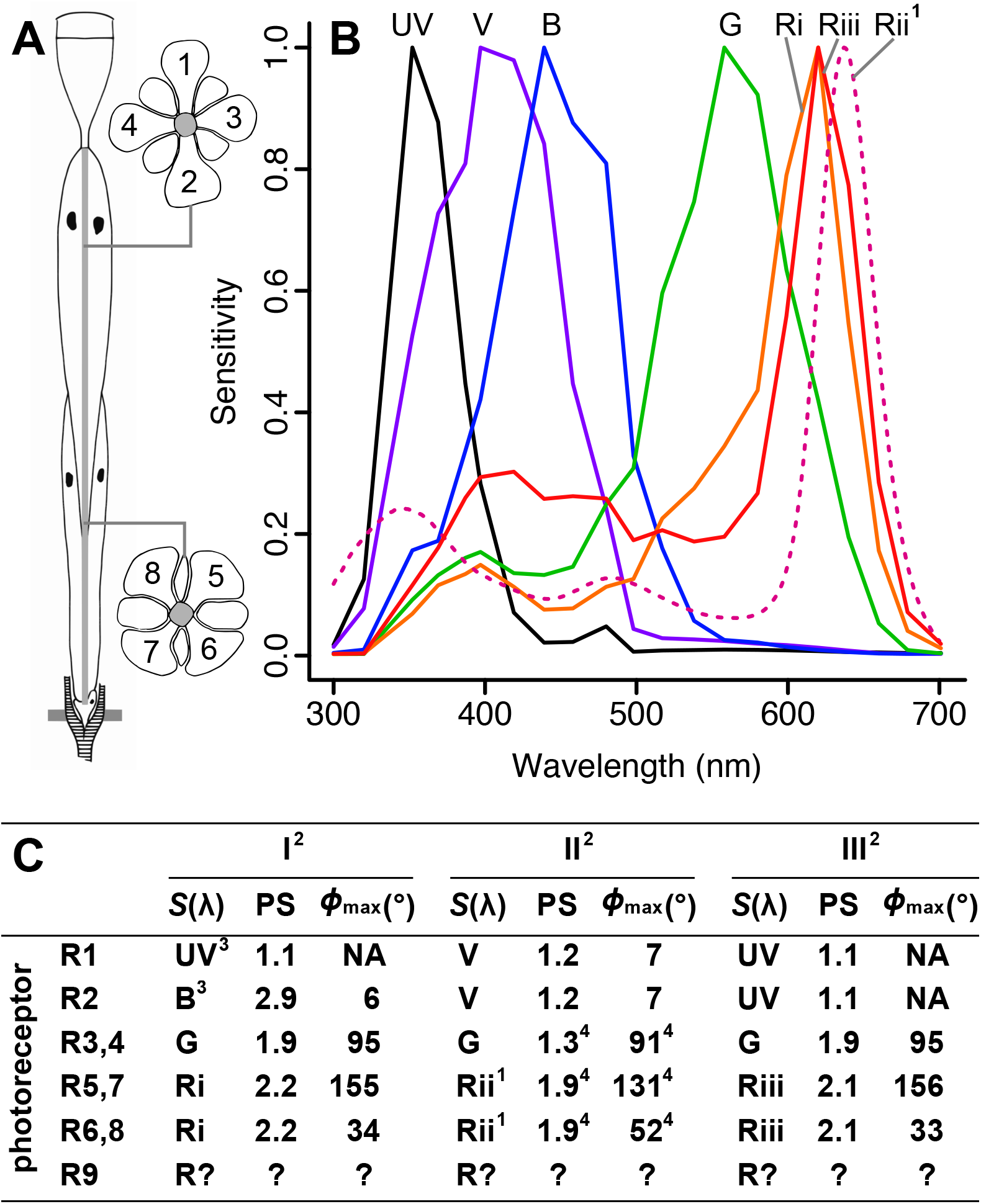
Visual system of female *Pieris rapae*. **A**, Diagram of ommatidium showing the arrangement of the nine photoreceptors (R1-9). **B**, Spectral sensitivities, S(λ), of the various photoreceptor spectral classes. Ultraviolet (UV), violet (V), blue (B), green (G), type I-III red (Ri-Riii). ^1^Spectral sensitivity predicted from a model of the female ommatidium (Stavenga and Arikawa 2011). **C**, Table summarizing the spectral class and polarization characteristics (PS, *ϕ*_max_) of photoreceptors R1-9 in ^2^ommatidial types I-III. ^3^UV and blue photoreceptors are positioned opposite each other but are equally likely to be in the R1 or R2 position. ^4^Values inferred from electrophysiological recordings of male butterflies.

**Fig. 2.**
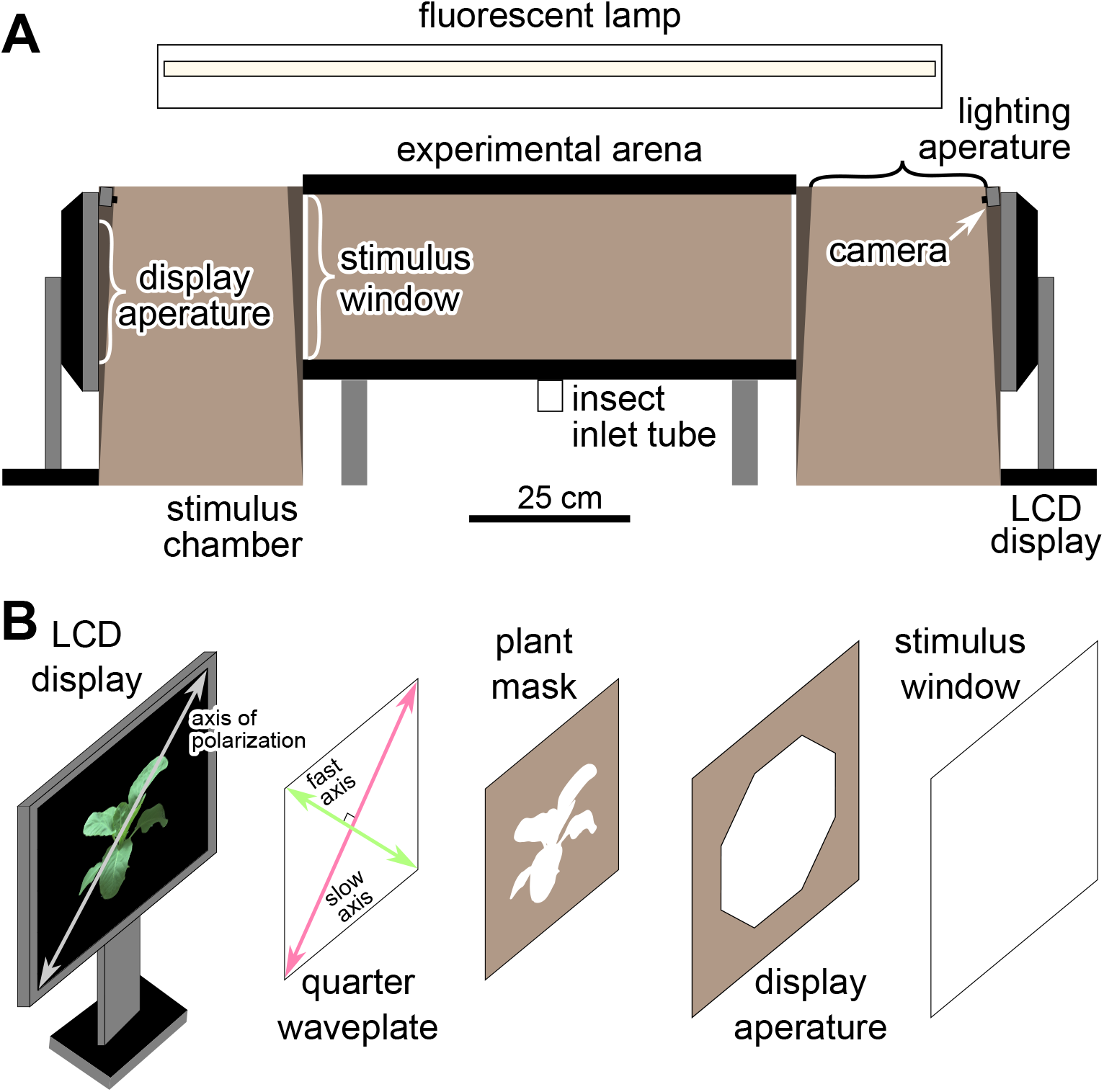
LCD monitor bioassay setup. **A**, Diagram of experimental arena. **B**, Exploded view of components between the LCD monitors and the stimulus windows. LCD = Liquid Crystal Display

The compound eye of *P. rapae* has been extensively characterized, but there is no obvious mechanism that would explain how *P. rapae* uses the signals from its suite of photoreceptors to discriminate among stimuli with different *DoLPs*. To determine whether *P. rapae* perceives differential *DoLPs* as differences in stimulus intensity or color, we sought to emulate the work of Kinoshita et al. (2011). In two-choice bioassays, we examined the responses of *P. rapae* to differences in *DoLP* or color between stimuli to determine whether intensity differences between the stimuli affected preference in a similar manner. We also determined the photoreceptors involved in *DoLP* discrimination by minimizing the blue, red, or blue and red light of cabbage images that we presented to *P. rapae* in bioassays. This type of manipulation is possible through use of our novel monitor bioassay (Blake et al., 2019b). We predicted that if a photoreceptor were involved in *DoLP* discrimination, then image manipulations of stimuli reducing the photoreceptor’s stimulation should alter the behavioral response of *P. rapae* to *DoLP* differences. We also modeled the catch of all *P. rapae* photoreceptors aiming to explain the observed behavioral bioassay responses of *P. rapae*.

## Methods

### Insect Material

Our laboratory colony of *P. r. rapae* originated from eggs obtained from the Carolina Biological Supply Company (# 144100, Burlington, NC, USA) and later from adults collected from cabbage fields near Delta, BC, Canada. Using a well-established protocol (Webb and Shelton, 1988), larvae were maintained on either a wheat-germ diet or on cabbage plants grown in a greenhouse. We housed both male and female adults in indoor cages (60 × 60 × 60 cm, BugDorm 2120, MegaView Science Co. Ltd., Taichung, Taiwan) kept at 18-25 °C and a photoperiod of 16L:8D. The females we tested in experiments were 3-14 days post eclosion and were assumed to be gravid. We tested females in multiple bioassays, each bioassay presenting a new pair of experimental plant images. These different bioassays were considered independent.

### General Experimental Setup

We used the same experimental arena (31.6 cm × 76.5 cm × 32.1 cm) and LCD monitor setup as recently described (Blake et al., 2019c; Fig. 2a). The inner surface of the removable arena lid was lined with matt white banner paper (NCR Corp., Duluth, GA, USA). We left the two end sections of the arena facing the monitors (stimulus windows) unobstructed but lined all the other inner surfaces of the arena with a matt brown kraft paper (NCR Corp.). To prevent build-up of any olfactory cues in the arena, we replaced the paper lining the interior surfaces and cleaned exposed glass surfaces with hexane daily.

In all experiments, we displayed cabbage plant images, created through photo polarimetry (Blake et al., 2019b), on paired liquid crystal display (LCD) monitors (1707FPt, Dell Inc., Round Rock, TX). These monitors were calibrated to minimize any differences between monitors in the displayed irradiance spectra of pixels with identical Red/Green/Blue (RGB) values (Fig. S1c). These monitors lack UV irradiance but the absence of UV wavelengths did not affect *DoLP-* based host plant preferences. We were able to independently manipulate both the *AoP* and *DoLP* of the plant image by rotating the monitor’s display and changing the alignment of the λ/4 retarder film (#88-253, Edmund Optics, Barrington, NJ) to the *AoP* of the display. Using LCD monitors also enabled us to readily manipulate both the plant image’s color and/or intensity.

The monitors were separated from the stimulus windows of the experimental arena by a stimulus chamber (31 cm × 31 cm × 47 cm) lined with the same kraft paper as the arena. This separation limited the range of viewing angles of the monitor from within the arena. In order to limit the visible portion of the LCD to that displaying the plant image, we placed a kraft paper plant mask over the display aperture in each stimulus chamber (Fig. 2b). The top of each stimulus chamber had a lighting aperture (27 × 26 cm) covered with the same white banner paper as the arena lid, thus affording similar illumination of the arena and the stimulus chambers. The arena and the chambers were lit from a florescent lamp (Fig. S2b; F32T8/SPX50/ECO GE, Boston, MA) centered 15 cm above the arena.

Using a camera mounted at the top rear of each stimulus chamber, we monitored the response of *P. rapae* females introduced into the arena. We allowed each female up to 5 min to approach one of the stimulus windows and recorded this approach as a behavioral response to the associated plant image. We considered females making no response non-responders. Image stimuli were alternated so they appeared equally often on both monitors/sides of the arena.

### Intensity-*vs*-Color Discrimination Experiment

To determine whether *P. rapae* females perceive differential *DoLP* as differential color or intensity, we performed experiments similar to those of Kinoshita et al. (2011). We presented females with paired stimuli consisting of the same cabbage image but modified to create differences in intensity (A), color and intensity (B), or *DoLP* and intensity between the two images (Fig. 3; Table S1). The paired stimuli we presented were (A) two unmodified images both with a *DoLP* of 31%; (B) one unmodified image and one red-shifted image each at a *DoLP* of 31%; and (C) two unmodified images presented with a *DoLP* of either 31% or 50%. In A-C, we presented the unmodified image with a *DoLP* of 31% at intensities lower (44%, 87%), equal (100%) and greater (130%) than the original intensity (Table S1). In (A), we did not present a choice between two unmodified images (*DoLP* 31%, 100% intensity) assuming no preference in response.

**Fig. 3.**
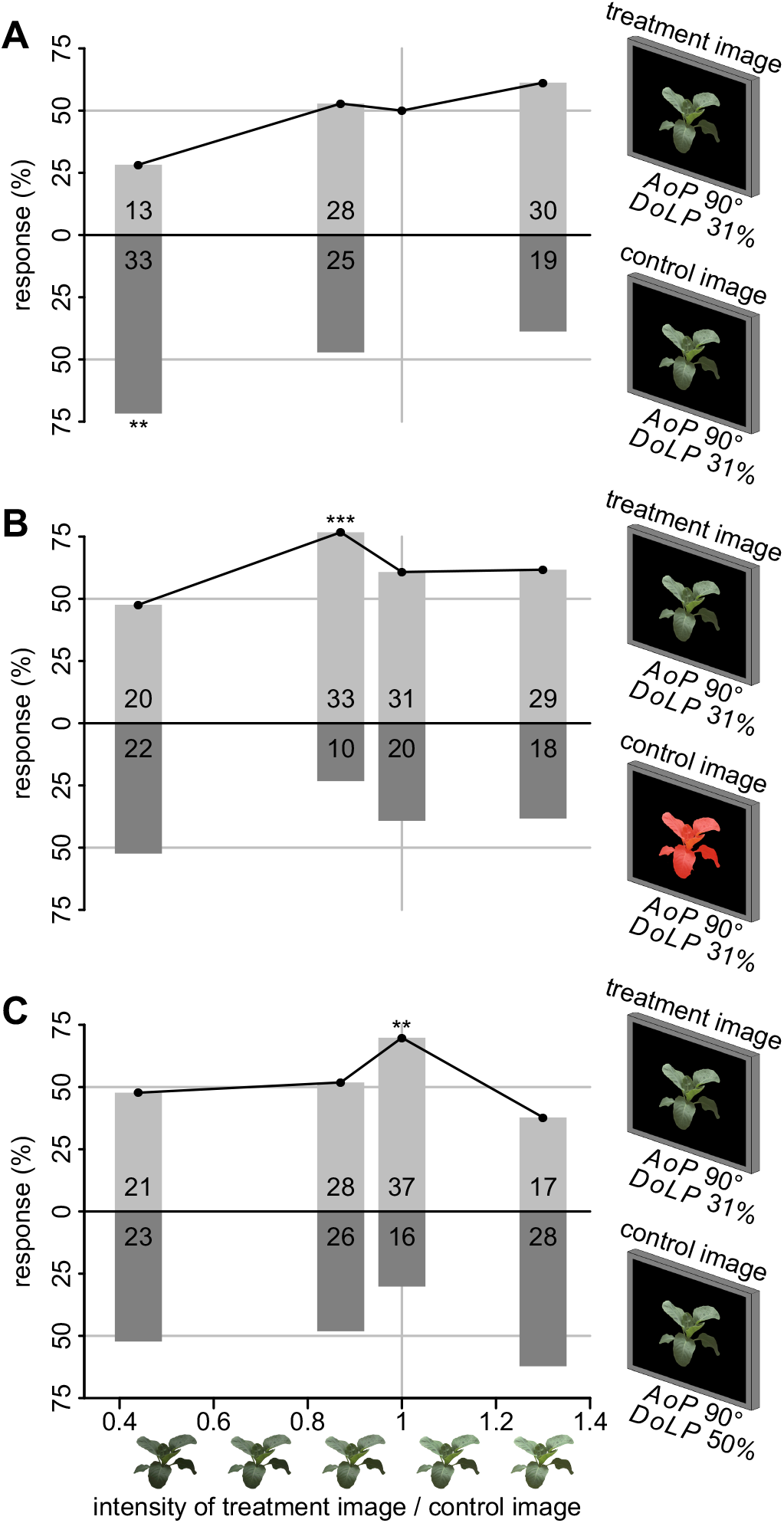
Intensity-*vs*-color discrimination experiment. Effect of relative increase in intensity of the treatment image on the preference of *Pieris rapae* females when treatment and control images differ only in intensity (**A**), in both color and intensity (**B**), and in both DoLP and intensity (**C**). The responses in (**A**) to treatment and control images of equal brightness were assumed to be 50%. Numbers of females responding to each stimulus are shown within bars. The asterisk(s) indicate(s) a proportion deviating from 50% (χ^2^ test, * *p* < 0.05, ** *p* < 0.01, \*\*\**p* < 0.001). *AoP* = Axis of Polarization; *DoLP* = Degree of Linear Polarization

### Color-Removal Experiment

To determine the photoreceptors involved in polarized light discrimination, we modified the color of cabbage images and offered *P. rapae* females a series of choices between these modified images presented at a *DoLP* of either 31% or 50%, with both images presented at an *AoP* of both 0° and 90°. To minimize the stimulation of the butterflies’ red photoreceptors, blue photoreceptors or both simultaneously (within the limits inherent in the RGB color space), we respectively set the red, blue, or red and blue values of all pixels in both stimulus images to 0 (Table S1). As a control, we also offered a choice between images with no modification to any pixel values.

### Statistical Analysis

We used chi-square tests to determine whether the proportion of *P. rapae* females responding to plant images differed from 0.5, and whether the proportion of females responding differed among the experimental treatments. We excluded non-responding females from statistical analyses.

### Modeling Photoreceptor Quantum Catches

Unless otherwise noted, all spectra were measured with a calibrated spectrophotometer (HR-4000, Ocean Optics Inc., Dunedin, FL, USA). To allow us to calculate the quantum catch of the background, we measured the ambient irradiance of the fluorescent lamps within the arena, the transmission of the arena wall and the λ/4 retarder film, and the reflectance of the brown kraft paper (Fig. S1a,b). We measured reflectance with a JAZ spectrometer (Ocean Optics) calibrated with a 99% Spectralon reflectance standard (SRS-99-010, Labsphere, NH, USA). Using photo polarimetry of the arena’s interior (Foster et al., 2018), we approximated the mean *DoLP* and modal *AoP* of the background across the human visible spectrum (400-700 nm) to be 10% and 90°, respectively.

We also used this spectrometer to measure the irradiance produced by the monitors at a range of 8-bit RGB values including pure red, green and blue spectra ([255, 0, 0], [0, 255, 0], [0, 0, 255], respectively, Fig. S1c) in order to estimate the monitor’s decoding gamma (*ɣ* = 1.90). Using equation 1 and the estimated decoding gamma *ɣ* we could appropriately scale and sum the pure spectra *I_C_*(λ) (where C is red, green or blue) using the red, green or blue pixel value *PV_C_* to estimate the displayed irradiance spectra across all wavelengths (λ) from 300 to 750 nm for any combination of RGB values. Using the mean RGB pixel values of the stimulus image, we could then create a mean spectrum for all pixels displayed in the image. The resulting spectrum was corrected for the transmission spectrum of the aquarium wall and the λ/4 retarder film (Fig. S1a).

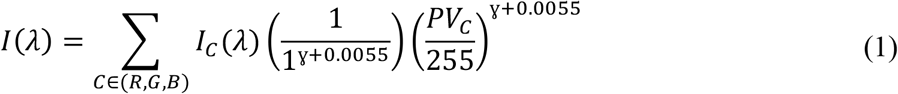

Using wavelength-specific effects of the λ/4 retarder film along with measurements of a photoreceptor’s *PS*, and the *AoP* of greatest sensitivity (*ϕ*_max_) taken from Blake et al. (2019a), we could use equation 2 to calculate the wavelength-specific effect of polarization on the photoreceptor’s response (*P_i_*(λ) for photoreceptor type *i*). This effect along with the previously mentioned intensity spectrum, and the reported spectral sensitivities of *P. rapae* photoreceptors (*R_i_*) (Blake et al., 2019a), allowed us to model the quantum catch (*Q_i_*) of all photoreceptor types with equation 3, with dλ being the spectral resolution of the spectrometer used to measure *I*(λ), and with all other variables interpolated to match this resolution. The quantum catch of the background (*Q_ib_*) was similarly calculated with irradiance *I*(λ) being determined from the irradiance spectra of fluorescence lamps and the reflectance spectra of the kraft paper. However, the measured values of *DoLP* and *AoP* (10% and 90°, respectively) determined from photo polarimetry were assumed to be uniform across 300-750 nm. As photoreceptors adapt to the background illumination, we then calculated the quantum catch relative to the background (*q_i_*) (equation 4).

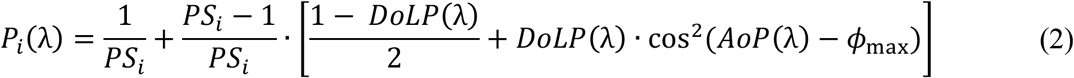

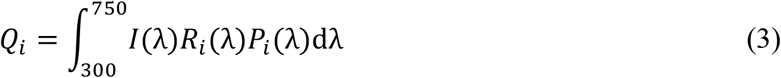

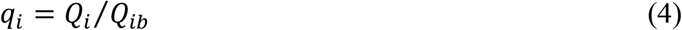

## Results

### Intensity-*vs*-Color Discrimination Experiment

In general, when treatment and control stimuli differed only in intensity, *P. rapae* females preferred the more intense stimulus (Fig. 3a). This preference was statistically significant only when the intensity of treatment stimuli was <50% of that of the control stimuli. When the treatment stimulus had an intensity of 87% relative to the control stimulus, females did not discriminate between these stimuli.

When we presented a choice between a red-shifted cabbage image and an unmodified treatment image of varying intensity, females significantly preferred the treatment image only with an intensity of 87% relative to the control image (Fig. 3b). Treatment images of a higher or a lower intensity were not significantly preferred, although there was a marginal preference for the treatment image when it had an intensity equal to, or greater than, that of the control image.

Similarly, when the treatment and control image differed in *DoLP*, females significantly preferred the treatment image only when it had an intensity equal to that of the control image (Fig. 3c). Treatment images of a lower intensity were as attractive as the control image while there was a non-significant preference for the control image when it was more intense.

### Color-Removal Experiment

When cabbage images were presented with all color channels intact (R+G+B), *P. rapae* females preferred the image with the lower *DoLP* both at an *AoP* of 0° and 90° (Fig. 4). When the blue color channel was removed (R+G), females shifted their preference towards the image with a higher *DoLP*, but only at an *AoP* of 0°. When the red color channel was removed (G+B), females preferred images with the higher *DoLP* at both *AoP*s. When only the green color channel of the image was included, females did not discriminate between images with a high or a low *DoLP*, when presented at an *AoP* of 90°. However, when these images were presented at an *AoP* of 0°, females chose the lower *DoLP* images similar to their response when all color channels were intact.

**Fig. 4.**
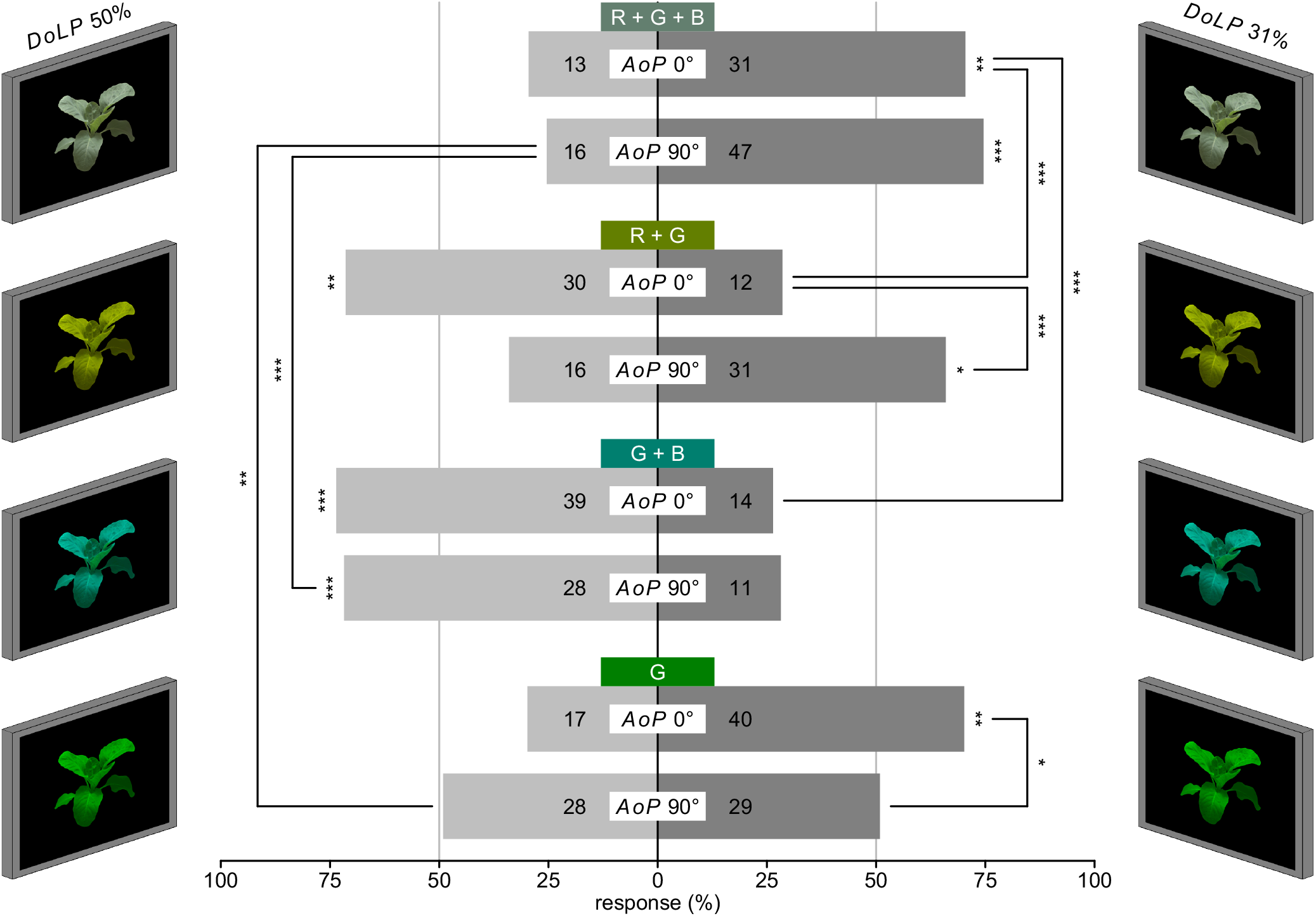
Color-removal experiment. Changes in the preference of *P. rapae* females for cabbage plant images differing in *DoLP*, with removal of RGB color channels. The stimulus images display unmodified RGB pixel values or have the red, blue, or red and blue values of all pixels in both stimulus images set to 0 (top to bottom). Numbers of females responding to each stimulus are shown within bars. The asterisk(s) indicate(s) a proportion deviating from 50% or a significant difference between two proportions (χ^2^ test, * *p* < 0.05, ** *p* < 0.01, *** *p* < 0.001). Note: the 31% *DoLP* is typical of cabbage plants. *AoP* = Axis of Polarization; *DoLP* = Degree of Linear Polarization

## Discussion

Our study refines the possible mechanisms for *DoLP*-based host plant discrimination by female *P. rapae*. According to our data, *P. rapae* females are likely not perceiving differences in *DoLP* as differences in purely intensity or in color. Rather, our data suggest that perception of color, intensity and polarization, at least in the context of host-plant discrimination, are all linked and contingent upon one another.

The intensity-*vs*-color discrimination experiment revealed that females preferred the plant image with greater intensity when all other factors were equal (Fig. 3). In our study, color preferences shifted in response to intensity changes in one of the two test stimuli, contrasting with results obtained in similar studies with *Papilio* butterflies (Kinoshita et al., 2011). While it is possible that *P. rapae* lacks true color vision (the ability to discriminate between colors independent of intensity), this explanation seems unlikely given the relatively close phylogenetic relationship between these butterflies and the similarities of their respective compound eyes (Kelber and Pfaff, 1999). Dissimilar experience of bioassay insects offers a more likely explanation for these contrasting results. While we tested the innate preferences of *P. rapae* females, corresponding studies with *Papilio* involved training (Kelber and Pfaff, 1999; Kinoshita et al., 2011). When paired images were similar in color and *DoLP*, we observed a positive linear relationship between the intensity of the treatment image and the preference of female *P. rapae* for this image (Fig. 3). In contrast, when image pairs were dissimilar in color or dissimilar in *DoLP*, female preference for the treatment image declined when the intensity of the treatment image was greater than that of the control image. Like in experiments with *Papilio*, these results suggest that differences in *DoLP* are being perceived as differences in color, albeit not independent of intensity.

The color-removal experiment revealed that blue, green and red photoreceptors are involved in the perception of differential *DoLP*. This conclusion is based on data showing (*i*) preferential responses to images with a lower *DoLP (AoP:* 0° and 90°) when all color channels were present; (*ii*) a preference shift for images (*AoP*: 0° or 90°) where either the blue or the red channel was removed; and (*iii*) the reversal of preferences with the green-only channel images (*AoP:* 90°) as compared with R+G or G+B images (*AoP*: 90°).

Contrary to results of the intensity-*vs*-color discrimination experiment, modeling of photoreceptor catch does not support the concept that differences in *DoLP* are perceived as color differences, at least not when modeled as a linear interaction among photoreceptors (Kelber, 2001; Kelber and Pfaff, 1999). The color triangles represent the modeled *P. rapae* color space and depict the relative quantum catch of the red, green and shortwave (omitting UV in type I) photoreceptors of the three ommatidial types disregarding intensity (Fig. 5,S2). In modeling the catch of the red photoreceptors, we assumed the catch of R5-8 are pooled negating much of *PS* of these photoreceptors. If *DoLP* discrimination could be explained through linear interactions between different photoreceptors, as seen in *Papilio* and in *P. rapae* with unpolarized stimuli, we would expect a consistent direction of preference between stimuli. For example, using existing linear color models for *Papilio* and *Pieris*, with the catch of green photoreceptors having a positive effect and blue and red receptors having a negative effect, we would expect the stimuli closest to the upper green vertex to be preferred. In our modeling, stimuli differing only in polarization characteristics largely align along the blue to green axis, with the direction of preference among paired stimuli tested converging on no one region of the color space (Fig. 5). This inconsistency applied to all ommatidial types (Fig. S2), albeit with smaller separations among low and high *DoLP* stimuli due to lower *PS* of the photoreceptors, and would likely not change even if photoreceptors were to be compared among different ommatidial types.

**Fig. 5.**
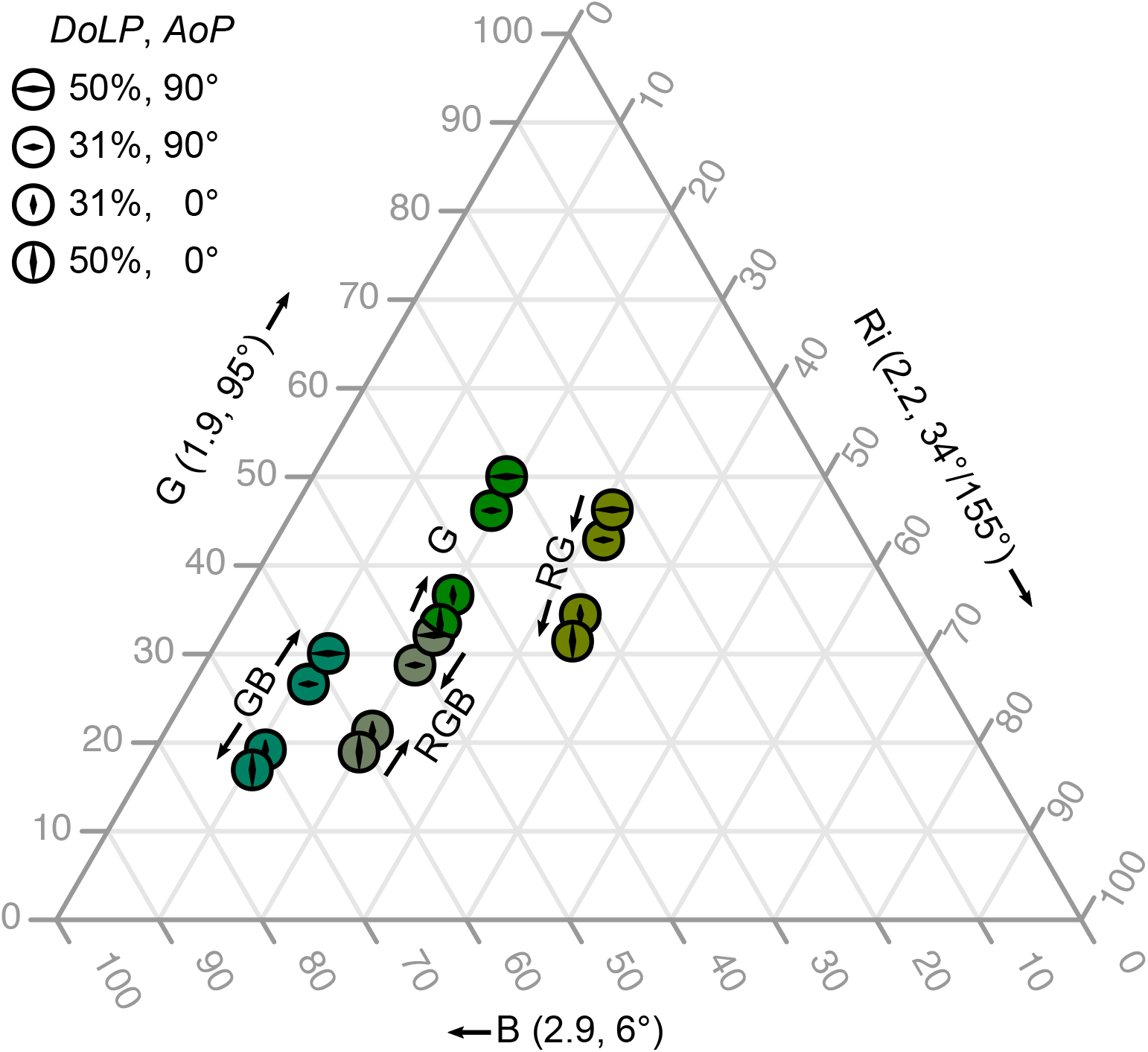
Color triangle representing the modeled color space of *Pieris rapae* females. This triangle shows a model of relative blue (B), green (G) and red photoreceptors’ (Ri) quantum catch in type I ommatidia. This color does not include the UV photoreceptor which was deemed acceptable due to the low levels of illumination in the UV and the low *PS* of UV photoreceptors. The numbers in parentheses show the *PS* and *ϕ*_max_ of each receptor. The colored circles show the stimuli in the color-removal experiment. Arrows indicate the stimuli preferred by female *P. rapae*.

Other plausible mechanisms also fail to explain our bioassay results. If polarization discrimination by *P. rapae* were to be dependent on comparisons between any two polarization-sensitive photoreceptors, or between one polarization-sensitive and one insensitive photoreceptor, we would expect *AoP* to have a strong effect on preference (Fig. S3a; How and Marshall, 2014), similar to how *Papilio* butterflies strongly prefer horizontally over vertically polarized light (Kelber, 2001). We would also expect such comparisons among photoreceptors to result in either a linear increase or decrease in preferential response as *DoLP* increased (Fig. S3b; How and Marshall, 2014). Yet, we found that the attractiveness of test stimuli was not affected by *AoP* outside regions near 45° and 135° and that images with a *DoLP* near 31% are preferred, with the appeal of stimuli declining both above and below this 31% value (Blake et al., 2019b). Comparisons between two or more pairs of photoreceptors are also unlikely to explain the observed *DoLP* preferences of *P. rapae* (Fig. S3ef). Models that incorporated the absolute value of the differences in responses between photoreceptors (Fig. S3cd; Meglič et al., 2019) could explain observed *AoP* preferences in *P. rapae*, but again would fail to explain *DoLP* preferences. The results of the color-removal experiment preclude true polarization vision (the ability to discriminate among stimuli independent of color or intensity), as changes in color prompted large shifts in polarization preference.

Our combined results suggest that a new and as of yet undescribed mechanism of polarization sensitivity underlies *DoLP* discrimination in *P. rapae*. The mechanism likely involves blue, green and red photoreceptor classes, and is affected by intensity, color and polarization. If true, this would be yet another example of unique neural processing of polarization information from object-reflected light. The systems for processing such information differ between all taxa thus far studied, including crabs (Smithers et al., 2019), fruit flies (Wernet et al., 2012), horse flies (Meglič et al., 2019), and backswimmers (Schwind, 1984). There are even as many as three different systems at work in *Papilio* butterflies depending on the behavioral context (Kelber et al., 2001; Kinoshita et al., 2011; Stewart et al., 2019). This information seems to show that different arthropod taxa have utilized the polarization sensitivity inherent in rhabdomeric photoreceptors to create visual subsystems tuned in accordance to their particular ecology. Further investigations into different arthropod taxa will almost certainly reveal novel combinations and processing of photoreceptor inputs using polarized light for object recognition.

## Acknowledgments

This study was supported by an Alexander Graham Bell Canadian Graduate Scholarship to AJB, Natural Sciences and Engineering Research Council of Canada (NSERC) - Undergraduate Student Research Awards to GSH, HG, SK and DM, and by an NSERC-Industrial Research Chair (IRC) to GG.

## Data Accessibility

Data are available from the Dryad Digital Repository https://doi.org/10.5061/dryad.xgxd254bs [32].

## Author Contributions

GSH, HG, SK and DM performed the bioassays. AJB and GG designed experiments. AJB preformed modeling and analysis. GG supervised the project. AJB wrote the first draft of the paper and GG provided comments.

---

**Table S1.**
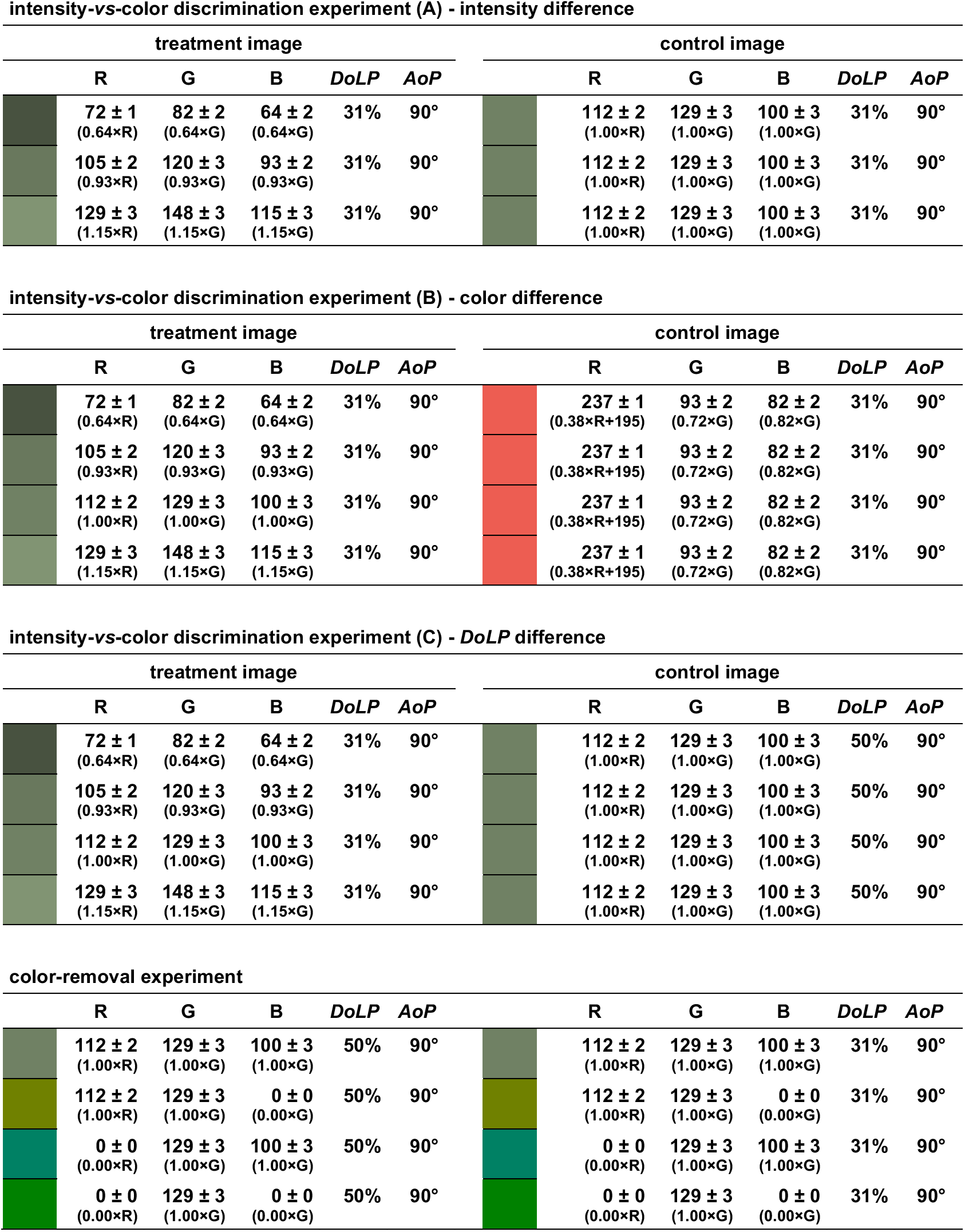
The mean RGB pixel values of plant images along with the corrections necessary to generate these images from the unmodified originals. The mean values were calculated from individual RGB means of each image. Also included are the degree of linear polarization (*DoLP*) and axis of polarization (*AoP*) of stimuli used in each experiment.

**intensity-vs-color discrimination experiment (A) - intensity difference**
| treatment image |  |  |  |  |  | control image |  |  |  |  |  |
| --- | --- | --- | --- | --- | --- | --- | --- | --- | --- | --- | --- |
|  | R | G | B | *DoLP* | *AoP* |  | R | G | B | *DoLP* | *AoP* |
|  | 72 ± 1<br>(0.64×R) | 82 ± 2<br>(0.64×G) | 64 ± 2<br>(0.64×G) | 31% | 90° |  | 112 ± 2<br>(1.00×R) | 129 ± 3<br>(1.00×G) | 100 ± 3<br>(1.00×G) | 31% | 90° |
|  | 105 ± 2<br>(0.93×R) | 120 ± 3<br>(0.93×G) | 93 ± 2<br>(0.93×G) | 31% | 90° |  | 112 ± 2<br>(1.00×R) | 129 ± 3<br>(1.00×G) | 100 ± 3<br>(1.00×G) | 31% | 90° |
|  | 129 ± 3<br>(1.15×R) | 148 ± 3<br>(1.15×G) | 115 ± 3<br>(1.15×G) | 31% | 90° |  | 112 ± 2<br>(1.00×R) | 129 ± 3<br>(1.00×G) | 100 ± 3<br>(1.00×G) | 31% | 90° |

**intensity-vs-color discrimination experiment (B) - color difference**
| treatment image |  |  |  |  |  | control image |  |  |  |  |  |
| --- | --- | --- | --- | --- | --- | --- | --- | --- | --- | --- | --- |
|  | R | G | B | *DoLP* | *AoP* |  | R | G | B | *DoLP* | *AoP* |
|  | 72 ± 1<br>(0.64×R) | 82 ± 2<br>(0.64×G) | 64 ± 2<br>(0.64×G) | 31% | 90° |  | 237 ± 1<br>(0.38×R+195) | 93 ± 2<br>(0.72×G) | 82 ± 2<br>(0.82×G) | 31% | 90° |
|  | 105 ± 2<br>(0.93×R) | 120 ± 3<br>(0.93×G) | 93 ± 2<br>(0.93×G) | 31% | 90° |  | 237 ± 1<br>(0.38×R+195) | 93 ± 2<br>(0.72×G) | 82 ± 2<br>(0.82×G) | 31% | 90° |
|  | 112 ± 2<br>(1.00×R) | 129 ± 3<br>(1.00×G) | 100 ± 3<br>(1.00×G) | 31% | 90° |  | 237 ± 1<br>(0.38×R+195) | 93 ± 2<br>(0.72×G) | 82 ± 2<br>(0.82×G) | 31% | 90° |
|  | 129 ± 3<br>(1.15×R) | 148 ± 3<br>(1.15×G) | 115 ± 3<br>(1.15×G) | 31% | 90° |  | 237 ± 1<br>(0.38×R+195) | 93 ± 2<br>(0.72×G) | 82 ± 2<br>(0.82×G) | 31% | 90° |

**intensity-vs-color discrimination experiment (C) - *DoLP* difference**
| treatment image |  |  |  |  |  | control image |  |  |  |  |  |
| --- | --- | --- | --- | --- | --- | --- | --- | --- | --- | --- | --- |
|  | R | G | B | *DoLP* | *AoP* |  | R | G | B | *DoLP* | *AoP* |
|  | 72 ± 1<br>(0.64×R) | 82 ± 2<br>(0.64×G) | 64 ± 2<br>(0.64×G) | 31% | 90° |  | 112 ± 2<br>(1.00×R) | 129 ± 3<br>(1.00×G) | 100 ± 3<br>(1.00×G) | 50% | 90° |
|  | 105 ± 2<br>(0.93×R) | 120 ± 3<br>(0.93×G) | 93 ± 2<br>(0.93×G) | 31% | 90° |  | 112 ± 2<br>(1.00×R) | 129 ± 3<br>(1.00×G) | 100 ± 3<br>(1.00×G) | 50% | 90° |
|  | 112 ± 2<br>(1.00×R) | 129 ± 3<br>(1.00×G) | 100 ± 3<br>(1.00×G) | 31% | 90° |  | 112 ± 2<br>(1.00×R) | 129 ± 3<br>(1.00×G) | 100 ± 3<br>(1.00×G) | 50% | 90° |
|  | 129 ± 3<br>(1.15×R) | 148 ± 3<br>(1.15×G) | 115 ± 3<br>(1.15×G) | 31% | 90° |  | 112 ± 2<br>(1.00×R) | 129 ± 3<br>(1.00×G) | 100 ± 3<br>(1.00×G) | 50% | 90° |

**color-removal experiment**
|  | R | G | B | *DoLP* | *AoP* |  | R | G | B | *DoLP* | *AoP* |
| --- | --- | --- | --- | --- | --- | --- | --- | --- | --- | --- | --- |
|  | 112 ± 2<br>(1.00×R) | 129 ± 3<br>(1.00×G) | 100 ± 3<br>(1.00×G) | 50% | 90° |  | 112 ± 2<br>(1.00×R) | 129 ± 3<br>(1.00×G) | 100 ± 3<br>(1.00×G) | 31% | 90° |
|  | 112 ± 2<br>(1.00×R) | 129 ± 3<br>(1.00×G) | 0 ± 0<br>(0.00×G) | 50% | 90° |  | 112 ± 2<br>(1.00×R) | 129 ± 3<br>(1.00×G) | 0 ± 0<br>(0.00×G) | 31% | 90° |
|  | 0 ± 0<br>(0.00×R) | 129 ± 3<br>(1.00×G) | 100 ± 3<br>(1.00×G) | 50% | 90° |  | 0 ± 0<br>(0.00×R) | 129 ± 3<br>(1.00×G) | 100 ± 3<br>(1.00×G) | 31% | 90° |
|  | 0 ± 0<br>(0.00×R) | 129 ± 3<br>(1.00×G) | 0 ± 0<br>(0.00×G) | 50% | 90° |  | 0 ± 0<br>(0.00×R) | 129 ± 3<br>(1.00×G) | 0 ± 0<br>(0.00×G) | 31% | 90° |

**Fig. S1.**
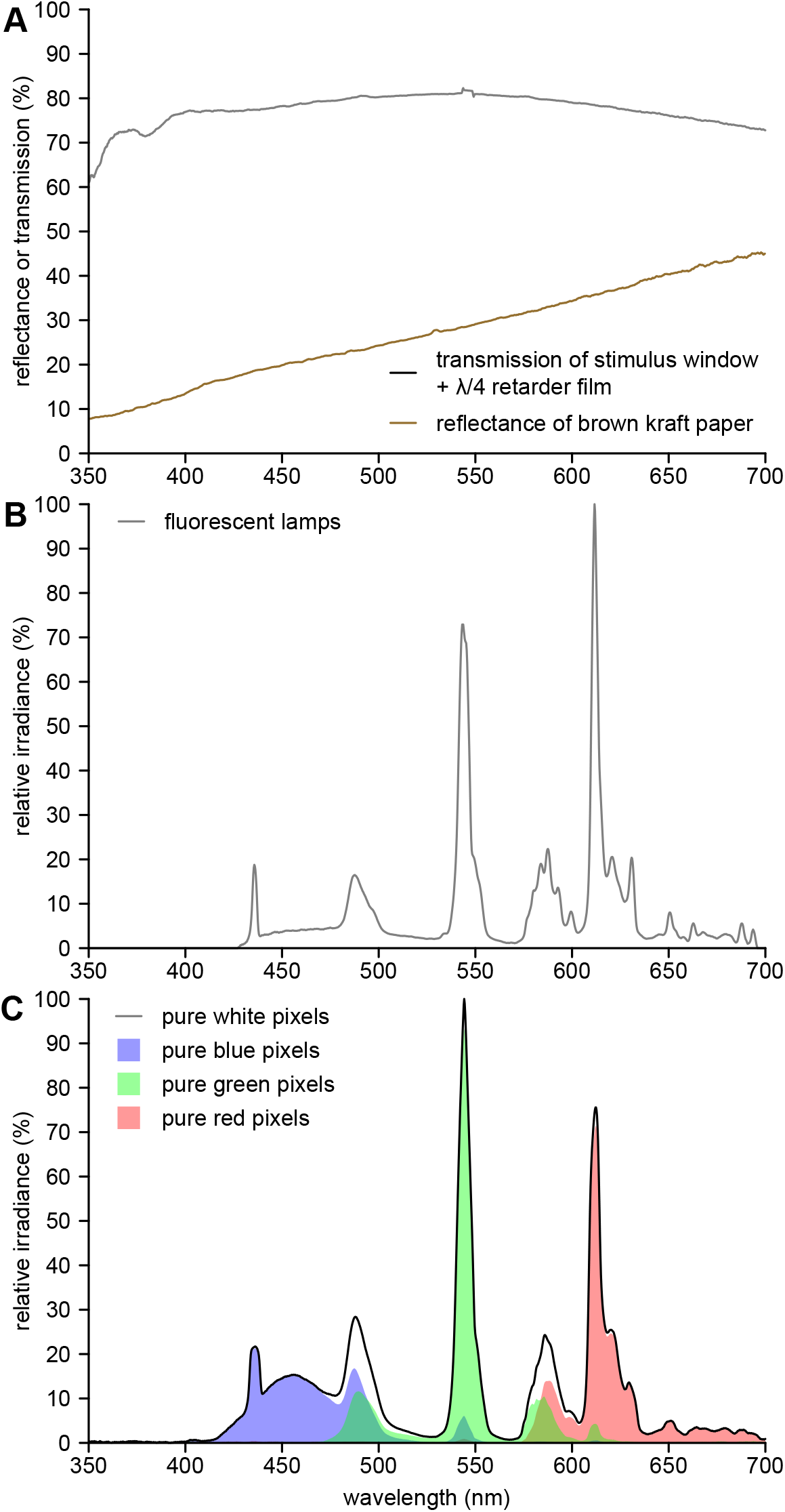
Spectra of filters, background, and illumination sources. **A**, Transmission spectrum of the stimulus windows of the experimental arena (Fig. 2) with a λ/4 retarder film and reflectance spectrum of the background brown kraft paper. **B**, Relative irradiance of the fluorescent lamps. **C**, Relative irradiance of white (RGB: 255, 255, 255), blue (0, 0, 255), green (0, 255, 0), or red pixels (0, 0, 255) as displayed on the bioassay monitors (mean of both LCD monitors).

**Fig. S2.**
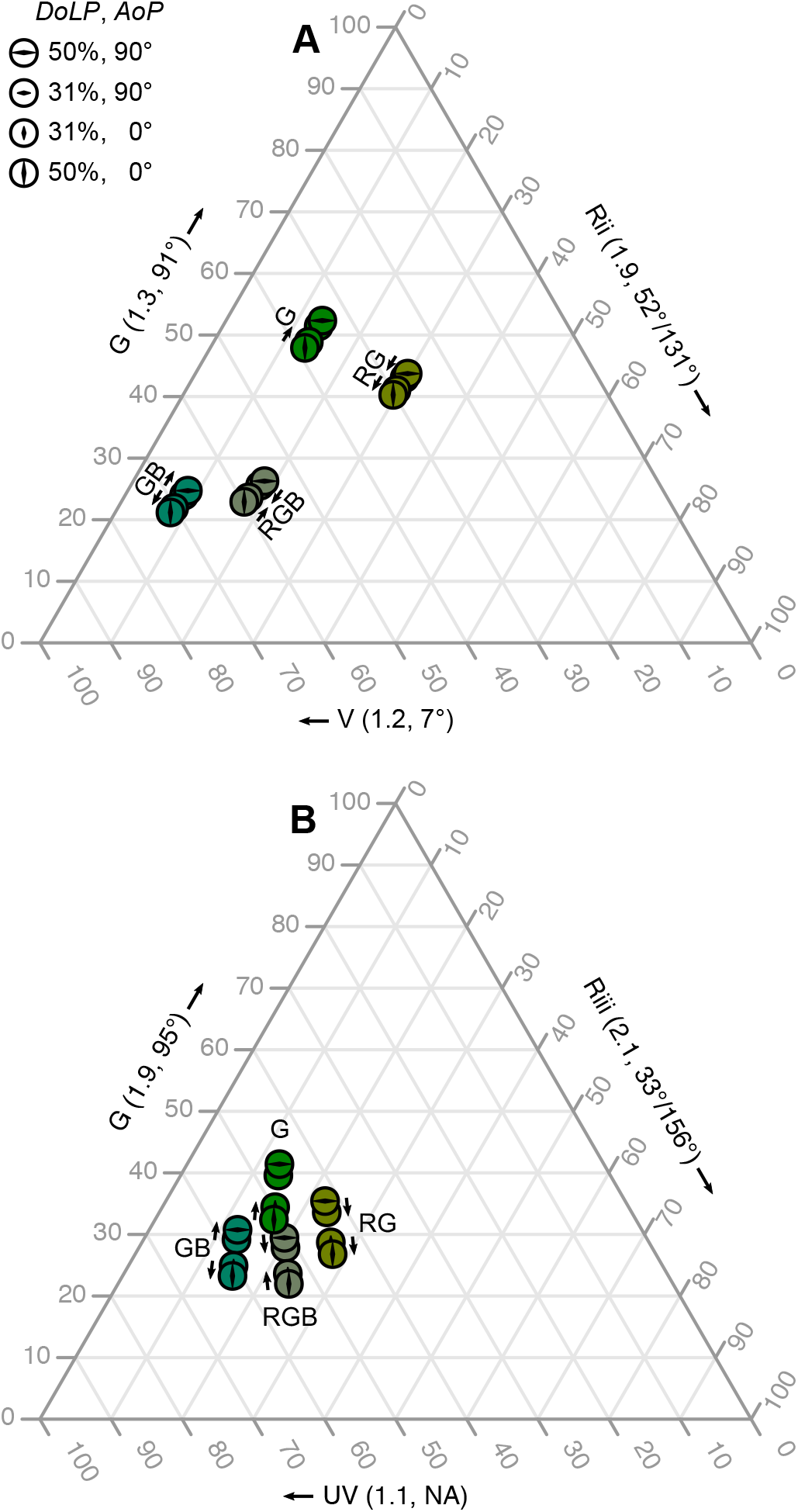
Color triangles representing the modeled color space of *Pieris rapae* females. Triangles show a model of relative ultraviolet (UV), violet (V), green (G) and red photoreceptors’ (Rii or Riii) quantum catch in type II and III ommatidia. Numbers in parentheses show the *PS* and *ϕ*_max_ of each receptor. The colored circles show the stimuli tested in the color-removal experiment. Arrows indicate the stimuli preferred by female *P. rapae*.

**Fig. S3.**
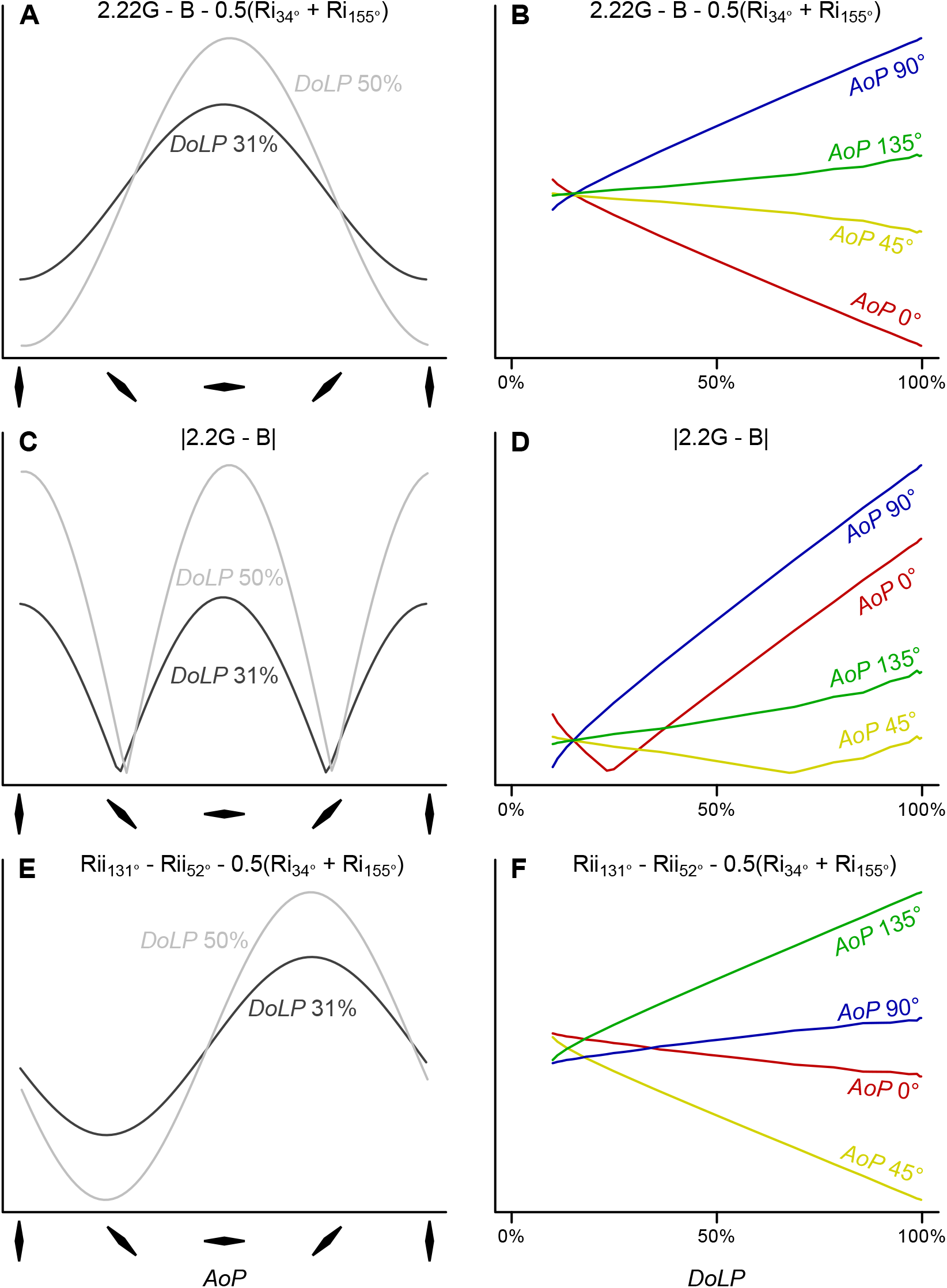
Effect of *AoP* and *DoLP* image manipulations on models combining photoreceptor catch from *Pieris rapae* females. (**A**,**C**,**E**), Effect of *AoP* on models at *DoLPs* of 31% and 50%. (**B**,**D**,**F**), Effect of *DoLP* on models at *AoP*s of 0°, 45°, 90°, and 135°. (**A**,**B**), Color model involving red, green and blue photoreceptors from type I ommatidia (Fig. 5) which would also be representative of any comparisons between polarization-sensitive photoreceptors, or between one polarization-sensitive and one polarization-insensitive photoreceptor. (**C**,**D**), Model calculating the absolute difference between two photoreceptors. (**E**,**F**), Model comparing more than two photoreceptors. *AoP* = Axis of Polarization; *DoLP* = Degree of Linear Polarization

